# Automated search of stimulation targets with closed-loop transcranial magnetic stimulation

**DOI:** 10.1101/2020.03.05.978445

**Authors:** Aino E. Tervo, Johanna Metsomaa, Jaakko O. Nieminen, Jukka Sarvas, Risto J. Ilmoniemi

## Abstract

Transcranial magnetic stimulation (TMS) protocols often include a manual search of an optimal location and orientation of the coil or peak stimulating electric field to elicit motor responses in a target muscle. This target search is laborious, and the result is user-dependent. Here, we present a closed-loop search method that utilizes automatic electronic adjustment of the stimulation based on the previous responses. The electronic adjustment is achieved by multi-locus TMS, and the adaptive guiding of the stimulation is based on the principles of Bayesian optimization to minimize the number of stimuli (and time) needed in the search. We compared our target-search method with other methods, such as systematic sampling in a predefined cortical grid. Validation experiments on five healthy volunteers and further offline simulations showed that our adaptively guided search method needs only a relatively small number of stimuli to provide outcomes with good accuracy and precision. The automated method enables fast and user-independent optimization of stimulation parameters in research and clinical applications of TMS.

## 1 Introduction

Neurons in the brain can be excited by transcranial magnetic stimulation (TMS). In TMS, a strong current pulse is fed into a coil placed on the scalp to induce an electric field (E-field) in the cortex (Barker *et al.* 1985). In addition to its use in neuroscience (Lisanby *et al.* 2000, Valero-Cabré *et al.* 2017), TMS is increasingly used in clinical applications ranging from preoperative mapping of motor and speech areas to treatment of various brain disorders such as depression and pain (see, *e.g.*, Lefaucheur and Picht 2016, Lefaucheur *et al.* 2014).

A typical TMS session starts with searching for the optimal stimulation parameters, such as location and orientation of the coil or the E-field maximum, to activate a muscle under investigation most effectively. These optimal stimulation parameters are often referred to as the motor hotspot and defined as the stimulation location and orientation that elicit the largest motor evoked potentials (MEP) measured by electromyography (EMG) (Rossini *et al.* 2015), but the shortest MEP latency or the lowest motor threshold can also be used in the definition (Rossini *et al.* 1994). The motor threshold (MT) is often defined as the stimulation intensity that produces an MEP exceeding a predefined amplitude with a probability of 50%. The E-field at the cortical motor hotspot due to stimulation with MT intensity may serve as a reference when adjusting the stimulation intensity at any cortical site. Therefore, the target search in the motor cortex is often performed even when the actual stimulation site is outside the primary motor cortex.

Optimal stimulation parameters are often searched for by measuring a collection of MEPs when stimulating the cortex around the expected motor representation area, with manually varied stimulation parameters: the location, intensity, and direction of the maximum E-field. The challenge is that, even with fixed stimulation parameters, the MEP amplitude is a random variable, *i.e.*, it varies significantly from stimulation to stimulation due to, for example, excitability fluctuations along the corticospinal tract (Kiers *et al.* 1993). Therefore, target search is laborious and time-consuming. The target search involves also subjective decision making based on the operator’s experience, which makes its accuracy and repeatability (over operators) questionable. Sometimes the target may reside in an unexpected location (Ahdab *et al.* 2016, Bulubas *et al.* 2016), in which case the operator’s expert opinion may lead to biased localization results.

To make the target search less user-dependent, Meincke *et al.* (2016) and Harquel *et al.* (2017) automated the process with a robotically controlled TMS coupled in a closed loop with EMG feedback. Meincke *et al.* (2016) defined the target as the stimulation coil location leading to the lowest MT; their algorithm started with finding an MEP-positive area within a search grid and continued with evaluating the MT at each MEP-positive site. They reported millimeter-scale repeatability in target localization with an automated protocol that took approximately an hour to complete. Harquel *et al.* (2017) developed an algorithm to find the location maximizing the MEP amplitude using a probabilistic Bayesian approach in which MEP amplitudes were modeled with a 2-dimensional Gaussian function and choosing stimulation sites based on the estimated decrease in entropy. However, both algorithms require a large number of MEPs and are developed to search for the target location with a fixed stimulation orientation (45° from the midsagittal line). The optimal orientation has been shown to differ between hand muscles (Bashir *et al.* 2013) and may also differ by tens of degrees across individuals (Balslev *et al.* 2007). Therefore, in addition to the stimulation location, the stimulation orientation should be optimized in the target search. Moreover, the performance of both of the above-mentioned algorithms was tested by varying the coil location in a grid with a 7-mm spacing. Such a grid is relatively sparse, and it might be beneficial to carry out the target search in a denser grid.

To overcome these limitations, we developed an algorithm we named BOOST (Bayesian Optimization Of Stimulation Targeting) to automatically find the optimal stimulation parameters, location and orientation, eliciting the largest MEPs. The automated search with BOOST is an iterative process, in which the search result, *i.e.*, the estimate of the optimal target, is updated in each iteration based on the already collected MEPs. Due to the probabilistic nature of the target search, we approach it with Bayesian optimization (see, *e.g.*, Shahriari *et al.* 2016). More specifically, we apply Gaussian process regression (Rasmussen and Williams 2006) to model the MEP responses as a function of the stimulation location and orientation. An advantage of this model is that we make no strong assumptions about the shape of the MEP response function. We present two versions of the BOOST algorithm: (1) in KG-BOOST, the stimulation is adaptively guided with a knowledge-gradient method (Frazier *et al.* 2009, Scott *et al.* 2011, Frazier and Wang 2016), which efficiently optimizes noisy functions (Picheny *et al.* 2013); (2) in Grid-BOOST, MEPs are sampled in a predetermined grid without adaptive guiding. KG-BOOST is the method that we recommend, whereas Grid-BOOST was implemented for comparison.

We evaluated the performance of different versions of the BOOST algorithm in one-dimensional cases with multi-locus TMS (mTMS), which allows electronic adjustment of the stimulation location and orientation without moving the transducer (Koponen *et al.* 2018a, Nieminen *et al.* 2019), making closed-loop stimulation fast end effortless. We hypothesized that, to achieve a given level of performance in the target search, intelligent sampling with knowledge gradient (KG-BOOST) requires less samples than systematic stimulation (Grid-BOOST) in an evenly spaced grid. We also expected KG-BOOST to perform better than Grid-BOOST when the same number of stimuli are used.

## 2 Methods

In this section, we describe the BOOST algorithm for an automated search for the optimal TMS parameters. In addition, we present the experimental set-up and the measurement protocol used to test the performance of the algorithm.

### 2.1 Algorithm

First, we present the BOOST algorithm and underlying mathematical models in a general form that allows the search of TMS targets in a multidimensional space. Then, we present a one-dimensional version of the algorithm in detail and the model parameters we used to validate it.

#### 2.1.1 Bayesian optimization with Gaussian processes

Our algorithm is designed to find stimulation parameters that produce the largest motor responses. We model the *n*th MEP amplitude as

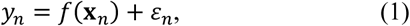

where *f*(**x**) is an unknown MEP response function that depends on *D* variables **x** = [*x*_1_, …, *x*_*D*_]^T^. Here, **x** can include, for example, the estimated location and orientation of the E-field maximum in the cortex. *ε*_*n*_ represents additive noise and is assumed to follow the normal distribution as 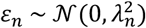, with variance 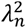 describing the variability of the MEP responses. Furthermore, *ε*_*n*_ for different *n* are assumed to be statistically independent of each other and, for simplicity, also of **x**_*n*_. The measured MEP response, *y*_*n*_, can be considered to be a random variable, *f*(**x**_*n*_) being the mean or expected value of this noisy sample of *f* at **x**_*n*_. Mathematically, the problem of finding the maximum of the MEP response function can be formulated as

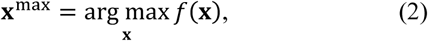

where **x**^max^ contains the stimulation parameters that maximize *f*. We assume that *f*(**x**) is a smooth function that indicates the effect of the E-field distribution on the neuronal pool responsible for the muscle contraction. The non-zero part of *f* informs us about the motor map of the investigated muscle.

The algorithm is based on modeling *f* with Gaussian process regression (Rasmussen and Williams 2006) given *N* noisy MEP responses **y** = [*y*_1_, …, *y*_*N*_]^T^. For stimulation parameters **X** = [**x**_1_, …, **x**_*N*_], we assume that the unknown values of *f* are jointly Gaussian

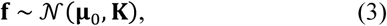

where **f** = [*f*(**x**_1_), …, *f*(**x**_*N*_)]^T^, **μ**_0_ = [*μ*_0_(**x**_1_), …, *μ*_0_(**x**_*N*_)]^T^ is a prior mean vector, and the covariance matrix **k** contains information on the *a priori* correlation of *f*(**x**_*n*_) and *f*(**x**_*m*_). In this context, we use a squared exponential covariance kernel defined as

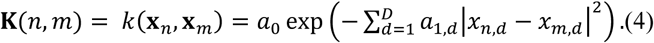

The model parameter *a*_0_ determines how much *f* can vary from the prior mean *μ*_0_, as one can see by setting *m* = *n* and observing that *k*(**x**_*n*_, **x**_*n*_) = *a*_0_ is the variance of *f*(**x**_*n*_). The second set of model parameters *a*_1,*d*_ (*d* = 1, …, *D*) determines the smoothness of *f, i.e.*, how quick changes there can be in each dimension *d*. The exponential term in Eq. (4) ensures that the correlation of *f*(**x**_*n*_) and *f*(**x**_*m*_) is large when the stimulation parameters **x**_*n*_ and **x**_*m*_ (with the elements *x*_*n,d*_ and *x*_*m,d*_, respectively) are close to each other and *a*_1,*d*_ are small. On the other hand, as *a*_1,*d*_ get large, **k** tends to a diagonal matrix, the samples *f*(**x**_*n*_) becoming mutually less correlated.

With the help of the described model and Bayesian inference, one can formulate the posterior probability distribution of *f*(**x**) at any point **x** given the measured *N* responses in vector **y** and the corresponding stimulation parameters **X**:

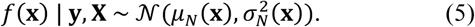

In this formula, the so-called posterior mean *μ*_*N*_(**x**) can be calculated as

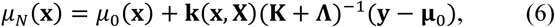

where **k**(**x, X**) = [*k*(**x, x**_1_), …, *k*(**x, x**_*N*_)] contains the covariances between **x** and **x**_1_, …, **x**_*N*_ similarly to Eq. (4), and **Λ** is a diagonal matrix with 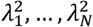 on its diagonal. The location of the posterior mean maximum 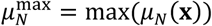 provides an estimate of the optimal stimulation parameters **x**^max^. One can also estimate the variance of 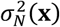 in Eq. (5):

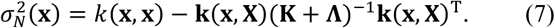

The sampling in the optimization process can be performed systematically in a grid of stimulation parameters as we do in the Grid-BOOST version of the algorithm. Another option is to utilize so-called acquisition functions that provide suggestions for the next sampling parameters **x**_*N*+1_ and help finding the optimum with a smaller number of samples than with the pre-determined grid approach. For this purpose, we apply the knowledge gradient (KG) that guides KG-BOOST optimization and is computed as

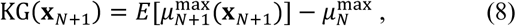

where 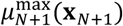 represents the maximum of the posterior mean function if we get one extra sample *y*_*N*+1_ corresponding to the parameters **x**_*N*+1_. In Eq. (8), 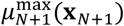 is a random variable depending on the random variable *y*_*N*+1_ and *E* stands for the mean operator. KG(**x**_*N*+1_) thus tells us how much the maximum of the posterior mean is expected to change if a new sample were collected with parameters **x**_*N*+1_. The optimally chosen next stimulation parameters lie where KG(**x**_*N*+1_) reaches its maximum. More details about the knowledge-gradient computation can be found in the Appendix. Note that there are also other methods to guide adaptive sampling, but we chose the knowledge-gradient method due to its reported good performance in optimizing noisy functions (Picheny *et al.* 2013).

#### 2.1.2 Algorithm for finding an optimal stimulation target in one dimension

The algorithm presented above works in multiple dimensions; here, we describe its one-dimensional version to find automatically either the optimal location of the peak E-field on a line segment or the optimal E-field orientation. Figure 1A depicts the flowchart of the algorithm. Several choices for the algorithm, such as the method for choosing the first stimulation parameters, were made intuitively. Most of these choices are justified by the fact that they resulted in a well-working algorithm. The automated search begins with determining the stimulation parameters for two initial samples within the search space [−*L*/2, *L*/2] that covers the selected range of stimulation locations or orientations (Step 1). The first initial sample is randomly chosen within the first half of the search space, *i.e., x*_1_ ∈ [−*L*/2, 0]. The second one is taken atlocation *x*_2_ = *x*_1_ + *L*/2. Next, a TMS pulse (or two pulses directly after Step 1) is given according to the determined stimulation parameter (Step 2), and the resulting MEP is measured from the target muscle (Step 3). For our analysis, we define the MEP amplitude *y*_*n*_ as the base-10 logarithm of the measured peak-to-peak amplitude (in microvolts), to meet better the Gaussian assumption of the MEP variability and to suppress large outlier MEPs, which could otherwise lead to an erroneous final result.

**Figure 1.**
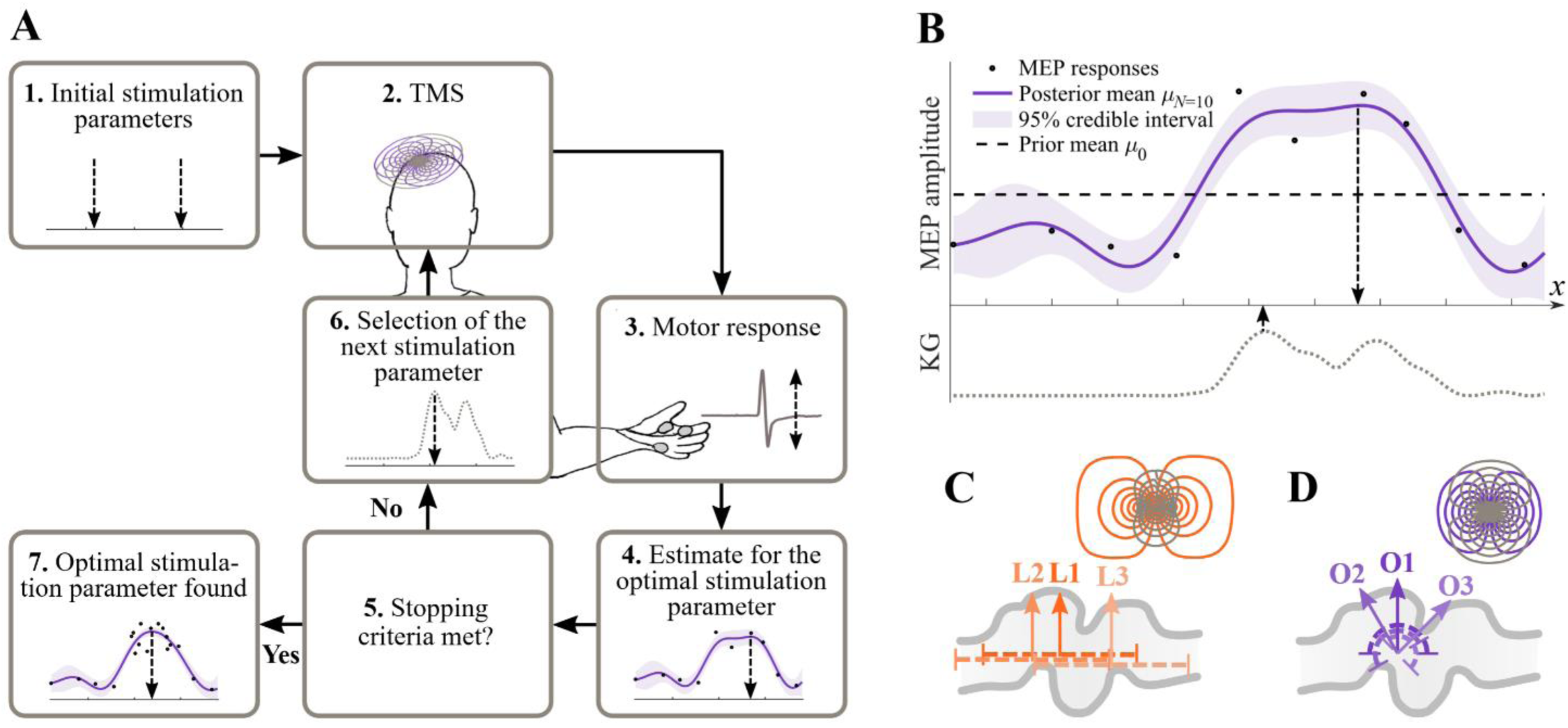
Bayesian optimization of stimulation targeting. A: Algorithm flowchart. B: Example of the estimation of the optimal stimulation target after 10 MEP responses (black dots). The purple solid line shows the posterior mean curve, and the shaded area depicts its 95% credible interval. The current estimate of the optimal stimulation target is where this curve reaches its maximum (downward-facing arrow). The black dashed line, being the mean of the posterior mean curve, depicts the prior mean for the next iteration. The next stimulus in KG-BOOST would be given with the parameter that maximizes the knowledge-gradient curve (gray dotted line; upward-facing arrow). C: Three placements (L1–L3) of the translation transducer over the primary motor cortex for 1D location search. The coil windings of the transducer are visualized in the top-right corner. D: The transducer placements for the orientation search (O1–O3) and the coil windings of the rotation transducer. In C and D, the dashed lines show the search space and the arrows indicate the location and orientation of the E-field maximum in the reference origin for each transducer placement. In each case, the stimulation of the reference origin is realized with the lower coil (orange or purple), and the stimulation of the other targets is achieved by feeding suitable current combinations to both of the overlapping coils.

In Step 4, we compute the estimate for the posterior mean with Eq. (6). Since the MEP variability differs between individuals, we adjust most of the model parameters adaptively based on the data gathered in the optimization process. As a prior mean for the first iteration, *i.e.*, after the two initial samples, we use a constant function, the output value of which is the average of the two initial responses. After the first iteration, the prior mean is still constant but now with an output value that is the average of the current posterior mean curve: 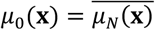, where 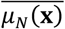 denotes the mean of the posterior mean curve (see the black dashed line in Fig. 1B). In the first iteration, we set the variability parameter as 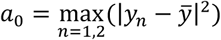, where 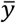 denotes the mean of the gathered MEPs. If the two initial responses happen to have the same value, we set *a*_0_ = (log_10_5)^2^ = 0.49. In the subsequent iterations, we define 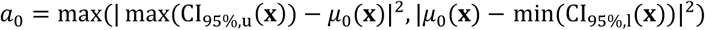 where 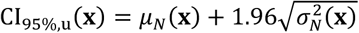 and 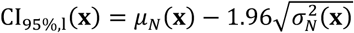 are the upper and lower limits of the 95% Bayesian credible interval of the posterior mean estimate, respectively (see an example of the credible interval in Fig. 1B). We define the smoothness parameter as *a*_1_ = *k*^2^π^2^/ (2*L*^2^), where *k* tells how many times *f* is expected to cross its mean value within the search space (see Rasmussen and Williams 2006, page 81). Here, we assume that *k* = 2. The MEP variance 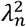 is chosen to be constant everywhere. In the first iteration, 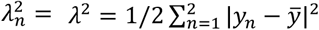, which is the variance of the elements in **y**. In the following iterations, 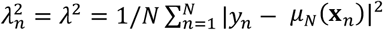, *i.e.*, the variance of the differences between the measured MEPs and the posterior mean curve. In each iteration, the estimate of the optimal stimulation target is where the posterior mean reaches its maximum.

In Step 5, the algorithm checks the stopping criteria. If they are not met, the next sampling point is determined (Step 6). The KG-BOOST algorithm stops if at least 14 responses have been collected and if, during the past eight iterations, the estimate of the optimal stimulation target has changed no more than 2 mm or 12° for the location and orientation search, respectively, or when 30 responses have been collected. These stopping criteria were formed based on preliminary convergence evaluations on test data. If the stopping criteria are not met, the next sampling point is chosen by evaluating the knowledge-gradient function (Eq. (8)) and by finding its maximum. If the knowledge gradient has the same value for all sampling points, we randomly pick the next stimulation parameter from 2–5 mm or 12–30° distance from the current estimate of the optimal stimulation target. We repeat Steps 2–6 until the stopping criteria are met. In Grid-BOOST, we take one sample in each point of an equally spaced grid in random order with no adaptive stopping criteria, sampling until the whole search space is covered systematically.

### 2.2 Data acquisition

Five healthy subjects volunteered for the study (aged 26–35 years, two males). All subjects were right-handed according to the Edinburgh inventory (Oldfield 1971). Prior to the measurements, each subject signed an informed consent. The study was approved by the Coordinating Ethics Committee of the Helsinki University Hospital and was carried out in accordance with the Declaration of Helsinki. TMS was administered with two different transducers connected to our in-house-developed mTMS system (Koponen *et al.* 2018a). One of the transducers comprises a figure-of-eight coil and an overlapping oval coil (Fig. 1C; Koponen *et al.* 2018a). With this translation transducer, we could electronically shift the location of the calculated E-field maximum along a 30-mm-long line segment in the cortex. The E-field in the cortex was calculated using a spherical head model with an 85-mm radius, the cortex assumed to be at 15-mm depth from the head surface. We had 31 possible locations (symmetrically around the reference origin with 1-mm spacing) of the E-field maximum along this line segment. The other transducer, with two overlapping figure-of-eight coils (Fig. 1D), allows electronic adjustment of the orientation of the maximum E-field (de Oliveira e Souza 2018). We restricted the possible stimulation orientations in the spherical head model to be within a 180° interval centered around the reference origin, with neighboring orientations separated by 1°. Here, the reference origin (0 mm/0°) means the location or orientation of the maximum E-field resulting from the stimulation with only the lower of the two overlapping coils (Figs. 1C, D). Thus, the reference origin moves together with the transducer. The applied pulse waveforms were monophasic (60-µs rise time, 30-µs hold period; Koponen et al. 2018b) and the interstimulus interval was randomized between 4 and 6 s.

The position of the mTMS transducers and the head of the subject were tracked with a neuronavigation system (eXimia 3.2, Nexstim Plc, Finland). For image-based guiding, we had T1-weighted magnetic resonance images of each subject. When needed, the position of the transducer with respect to the subject’s head was kept fixed with the help of the neuronavigation system, which allowed stimulation only when the transducer location was within 2 mm and all rotation angles less than 2° from their target values.

The motor responses were measured with surface EMG integrated in the eXimia 3.2 system (500-Hz low-pass filtering, 3-kHz sampling frequency). The silver/silver-chloride surface electrodes (Ambu Neuroline 720, Ambu A/S, Denmark) were in a bipolar arrangement with the active electrode placed over the muscle belly of the right *first dorsal interosseus* (FDI) and the reference electrode on the second proximal phalange. The ground electrode was placed on the back of the hand. The eXimia system analyzed the evoked responses and displayed the peak-to-peak amplitudes and latencies of the MEPs in real time. To get the MEP data into our algorithms programmed with Matlab (The MathWorks, Inc., USA), we imported the video stream of the eXimia system to our control computer in real time with a USB video grabber (DVI2USB 3.0, Epiphan Systems Inc., Canada). From the video stream, we extracted the MEP-amplitude and MEP-latency values as reported by the eXimia system. We also analyzed the baseline EMG signal 5–200 ms before the TMS pulse for real-time rejection of responses with muscle preactivation. We accepted an MEP if its onset latency was 15–30 ms and if the baseline EMG signal was within ±10 μV (when determining the MT) or ±15 μV (when running the KG-BOOST or Grid-BOOST algorithms). If these conditions were not met, we repeated the stimulation with the same parameters until the MEP response was acceptable. Since the automatic MEP analyzer of the eXimia system sometimes missed small responses, giving just 0 μV as their amplitude, we replaced for data analysis each 0-µV MEP amplitude with a random amplitude drawn from a uniform distribution with an interval of 5–15 μV.

Each subject had two measurement sessions, conducted on different days. At the beginning of the first session, we manually located the optimal stimulation target of the FDI muscle with the figure-of-eight coil of the translation transducer. For this, we delivered TMS pulses to the left primary motor cortex, varying the target location with millimeter-level steps around the hand-knob area. The stimulation intensity was first about 70 V/m, then adjusted so that the maximal MEP amplitude would be approximately 1 mV. The ISI was about 5 s. After delivering several tens of pulses around the hand-knob region to outline the MEP-positive area, we visually evaluated the distribution of the MEP responses and selected one target approximately from the center of the area showing the largest MEPs. During the manual search, the estimated orientation of the peak E-field was kept perpendicular to the overall orientation of the precentral gyrus. Next, we determined the resting MT (rMT) for the FDI muscle with a maximum-likelihood method (Awiszus 2003), applying 20 pulses with different intensities while having 50 µV as a threshold for MEP acceptance. For the rest of the two sessions, the stimulation intensity was set to 110% rMT.

In the first session, we performed the automated search for the optimal stimulation location with the KG-BOOST and Grid-BOOST algorithms described in Section 2.1.2. We positioned the transducer in three different locations to test whether the algorithm can locate the optimal stimulation site regardless of its placement within the search space. The first placement corresponded to the manually found FDI target (placement L1), the second was ∼5 mm to the medial (placement L2) and the third ∼8 mm to the lateral direction (placement L3) from L1 (see Fig. 1C). The transducer was kept fixed at L1–L3 and the peak E-field was electronically adjusted to one of the 31 possible locations according to the algorithm. We repeated both KG-BOOST and Grid-BOOST seven times for each transducer placement (L1–L3), resulting in 21 repetitions per subject. The transducer placements and the utilized version of the algorithm were applied in a pseudorandom order. The experimenters were aware of the transducer placement as well as the applied algorithm version during the experiments. With both KG-BOOST and Grid-BOOST, the estimated posterior mean curve computed with Gaussian process regression (Eq. (6)) was calculated on the 30-mm line segment with a grid spacing of 0.25 mm.

In the second session, we conducted the automated search for the optimal stimulation orientation. We set three transducer placements as follows: the placement O1 corresponded to L1, the placement O2 was oriented ∼30° counterclockwise, and the placement O3 ∼45° clockwise with respect to O1 (see Fig. 1D). The sampling in Grid-BOOST consisted of 31 pulses with 6° steps ranging from −90° to 90° around the reference origin. In KG-BOOST, we had the same 180°-wide search space with the possible stimulation orientations separated by 1° steps. We repeated both KG-BOOST and Grid-BOOST seven times with transducer placements O1–O3, resulting in 21 repetitions for each subject. The posterior mean curve was computed with a grid spacing of 0.5°.

### 2.3 Data analysis

In this Section, we first explain how we simulated other search methods using the measured data. Then, we show how we evaluated the accuracy of different search methods by comparing the optimization results with the ground truth and how we determined the precision as the deviation in the search outcomes. In addition, we present details of the statistical testing comparing the performance of KG-BOOST with the other methods. The investigators were not blinded when analyzing the data.

#### 2.3.1 Other search methods

To complement the results obtained directly from our experiments, we simulated the performance of three other sampling methods using the MEPs collected in the KG-BOOST and Grid-BOOST searches, sampling from these data without replacement.

First, we conducted sparser grid sampling (sparse Grid-BOOST) with the data collected in the original Grid-BOOST searches (that we call dense Grid-BOOST from now on). In sparse Grid-BOOST, the number of samples collected was equal to that of the corresponding KG-BOOST search repetition (14–30 samples per search). We selected every other sample from the denser grid and sampled from this subset of the data in random order until the total number of samples was the same as in the KG-BOOST search. If needed, we took extra samples from the unused half of the data. Since the two data subsets could be constructed in two ways, with the first one including either even or odd indices, we randomized the order in which the two subsets were used. The posterior mean curve corresponding to the sparse Grid-BOOST sampling was computed in the same way as with KG-BOOST and dense Grid-BOOST, *i.e.*, with the parameters presented in Section 2.1.2.

To mimic the target search without any modeling of the MEP responses, we also evaluated a search strategy in which the estimate of the optimal stimulation target coincided with the location of the maximum MEP response. For this, we used the data sampled in the sparse Grid-BOOST searches. We refer to this search strategy as the maximum-MEP method.

For comparison, we simulated the previously reported AutoHS method by Harquel *et al.* (2017) to find the optimal stimulation site. We made three adjustments to the AutoHS method due to differences in the MEP-sampling schemes: (1) Our sampling grid spacing was 1-mm (cortical grid) as opposed to 7 mm (grid of coil locations on the scalp) used in the original study. (2) Our search space was one-dimensional instead of two-dimensional. (3) We allowed sampling at each stimulation site at most once, whereas in Harquel *et al.* (2017) the same location was targeted at most twice. Because AutoHS gathers five MEPs at each iteration at the selected stimulation target and because for some targets we had collected only seven MEPs, we run the method only once for each subject and transducer placement. When searching for the optimal stimulation location with AutoHS, the possible values for the maximum MEP amplitude, the Gaussian width, and the center point of the Gaussian were {100 μV, 300 μV, …, 3900 μV}, {2 mm, 4 mm, …, 20 mm}, and {−15 mm, −14.75 mm, …, 15 mm}, respectively. For defining the next stimulation target in each iteration, the grid spacing was 1 mm. The first stimulation target and the center point of the prior distribution were always set in the middle of the search space (the reference origin).

We also performed the search for the optimal stimulation orientation with AutoHS although such an application was not described in the original article. Indeed, the shape of the Gaussian function could be expected to model the MEP distribution as a function of the stimulation orientation, too. Here, the possible values for the maximum MEP amplitude, the Gaussian width, and the center point of the Gaussian were {100 μV, 300 μV, …, 3900 μV}, {12°, 24°, …, 120°}, and {−90°, −89.5°, …, 90°}, respectively. The spacing of the sampling grid was 6°. The prior for the optimal angle was centered around the reference origin as in the location search. The width of the prior distribution was 30°.

#### 2.3.2 Estimation of accuracy and precision

To estimate the bias in the results (*i.e.*, the accuracy) obtained with different search strategies, we first defined the best estimate for the optimal stimulation target (hereafter, the ground truth) for each transducer placement and subject. This was done by first pooling the data from the seven repetitions of the KG-BOOST and Grid-BOOST searches. Each data pool included at least 315 MEP responses from the same spatial/angular distribution. For each of the 30 cases (five subjects, two transducers, three transducer placements), we computed a median curve in a grid with a 1-mm/1° spacing using a sliding window that took into account the responses that were closer than 5 mm (location search) or 30° (orientation search) from the computation point. We defined the ground truth as the location of the maximum of the median curve. If several points of this curve had the same maximum value, we defined the ground truth as their mean location. When calculating the ground truth, we replaced the amplitudes of those MEPs that the eXimia system had originally identified as 0-µV MEPs with the peak-to-peak amplitude of the EMG signal in the time interval of 15–45 ms after the TMS pulse.

We computed the average location/orientation of the search results over the seven repetitions, separately for each search strategy (KG-BOOST, dense and sparse Grid-BOOST, the maximum-MEP method), and compared it with the corresponding ground truth. To determine the group-level accuracy, we computed the mean of these differences in 15 cases (3 transducer placements × 5 subjects). This accuracy measure tells us how close the average result was to the ground truth. With AutoHS, we had in each case only one simulated search result and used its difference from the ground truth when computing the mean accuracy.

To assess the precision (degree of scatter) of each search method, we computed the standard deviation of the corresponding seven final search results and averaged these standard deviations over the transducer placements and subjects. This precision measure describes the repeatability of the outcome of each search method.

#### 2.3.3 Statistical analysis

The difference between the precision/accuracy of KG-BOOST and the other search methods with the same number of search repetitions (*i.e.*, dense and sparse Grid-BOOST, the maximum-MEP method) was tested by permutation statistics as follows. Altogether, 30 test values (5 subjects × 3 transducer placements × 2 search methods under comparison) were randomly divided into two groups 1,000,000 times. The accuracy and precision for both the location and the orientation search were treated separately. For each permutation, we computed the mean value for both groups. We obtained a two-tailed *p*-value as the proportion of permutations for which the absolute value of the difference of means of the permuted groups exceeded the absolute value of the corresponding original difference of means between the sampling methods. The level of statistical significance was set at 0.01. After Bonferroni correction for multiple comparisons (12 comparisons), the corrected significance level was 0.00083.

## 3 Results

Results of the location and orientation searches are visualized in Fig. 2. Figure 2A shows how the search results for a representative subject with the three different transducer placements L1–L3 are distributed with respect to the ground truth. The three smaller graphs on the right side of Fig. 2A depict an example of a single run with each of the four search methods (KG-BOOST, dense and sparse Grid-BOOST, the maximum-MEP method). Figure 2B shows similar example results for the orientation search. Figures 2C and 2D illustrate the error distributions of the search outcomes, *i.e.*, how far the optimized parameters are from the ground truth, for each subject.

**Figure 2.**
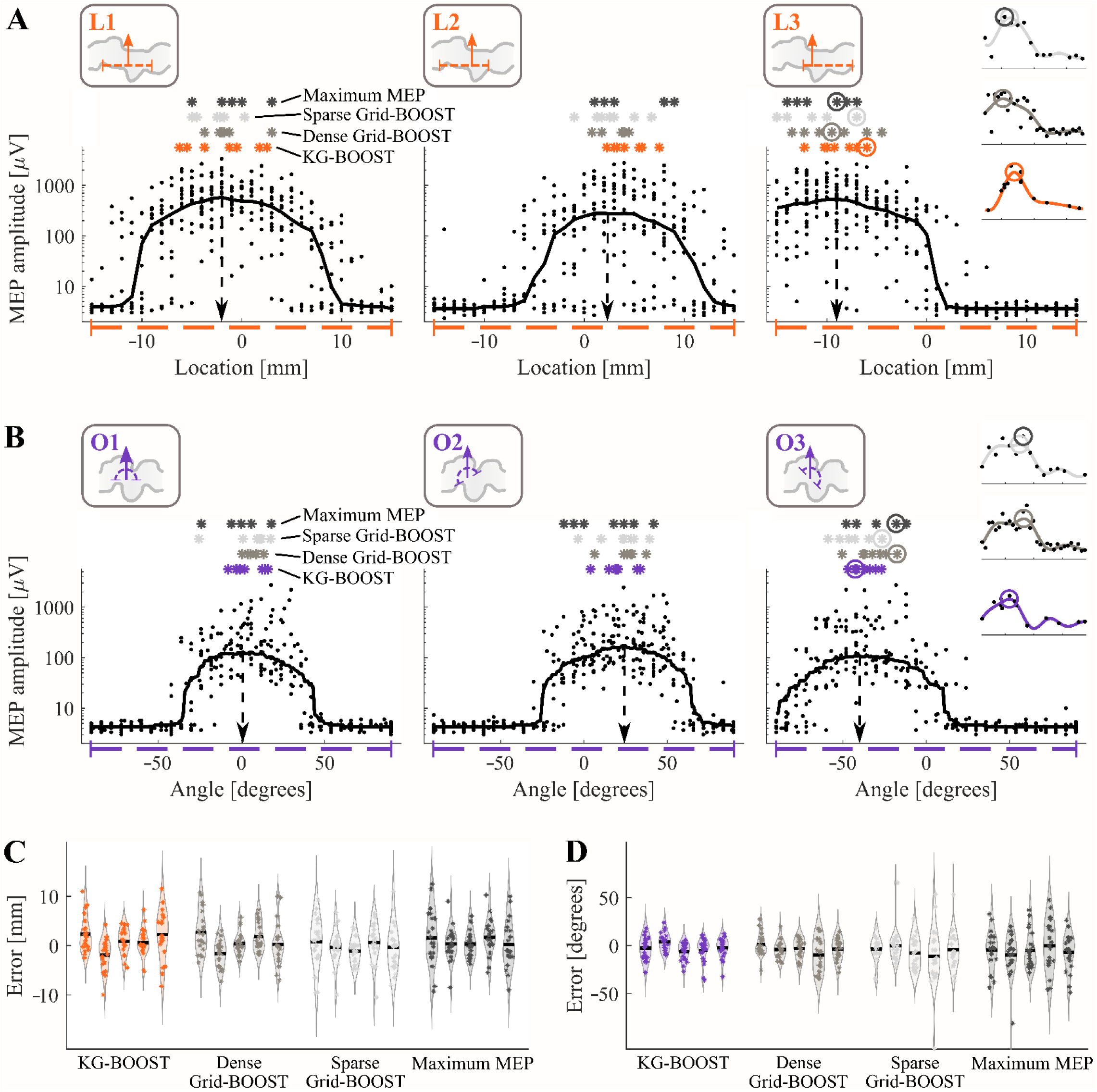
Results of the location and orientation searches. A: Location search with Subject 3 and transducer placements L1–L3. B: Orientation search with Subject 1 and transducer placements O1–O3. The transducer placements with respect to the hand-knob area in the pre-central gyrus are shown in the boxes on the top (the arrows indicate the location/orientation of the manually found target), with the corresponding results panels below them. The black dots show the MEP responses measured during all KG-BOOST and dense Grid-BOOST searches. The black line shows the moving median of these responses, and the location of its maximum (black dashed arrow) is the ground truth. The asterisks show the estimates of the optimal stimulation locations/orientations of the seven repetitions with KG-BOOST (orange/purple), dense (gray) and sparse (light gray) Grid-BOOST and the maximum-MEP method (dark gray), respectively. The three smaller graphs on the right visualize single runs of different search methods with L3/O3. In these graphs, the black dots illustrate the MEP responses, the solid lines depict the posterior mean curves, and the circles indicate the estimated optimal stimulation locations/orientations (the corresponding asterisks in the L3/O3 results panel are circled). C–D: Violin plots visualizing the error (difference from the ground truth) in the search of the optimal location (C) and orientation (D) for each subject, with the data of all transducer placements combined. The asterisks show the error of each search run, and the black lines indicate the mean errors.

The convergence of the adaptively guided KG-BOOST in the search for the optimal location and orientation is presented in Figs. 3A and 3B, respectively. Figure 3C depicts the performance metrics of the location search over subjects. The mean accuracy (*i.e.*, how far the average search result is from the ground truth) of KG-BOOST was 1.4 mm (range: 0.04–5.1 mm) which is similar to the mean accuracy of 1.5 mm (range: 0.2–5.0 mm) of dense Grid-BOOST (*p* = 0.87). The accuracies of sparse Grid-BOOST (mean: 2.0 mm; range: 0.04–4.4 mm) and the maximum-MEP method (mean: 2.1 mm; range: 0.3–5.3 mm) were slightly worse, although the differences were not statistically significant when compared to the accuracy of KG-BOOST (*p* = 0.32 for the sparse Grid-BOOST and *p* = 0.21 for the maximum-MEP method). The precision, expressing how repeatable the results are, was best with dense Grid-BOOST (mean: 2.7 mm; range: 1.2–5.2 mm). The mean precision of KG-BOOST (mean 3.2 mm; range 1.6– 5.3 mm), the maximum-MEP method (mean: 3.4 mm; range: 1.9– 6.6 mm) and sparse Grid-BOOST (mean: 3.4 mm; range: 1.7– 6.2 mm) were comparable to each other. When comparing KG-BOOST with dense or sparse Grid-BOOST or the maximum-MEP method, the *p*-values were 0.23, 0.68, and 0.72, respectively. With KG-BOOST, sparse Grid-BOOST and the maximum-MEP method, the average number of samples collected was 18 (range: 14–30), which means that on average the search took 1.5 min. With dense Grid-BOOST, we always gathered 31 samples per search, corresponding to an average time of 2.6 min. Moreover, the manually found optimal stimulation location, which corresponds to the reference origin with the transducer position L1, varied on average 3.5 mm (range: 0–6 mm) from the corresponding ground truth. The average number of stimuli in the manual search was 59 (range: 37–78), administered on average in 11 minutes (range: 4–18 min).

**Figure 3.**
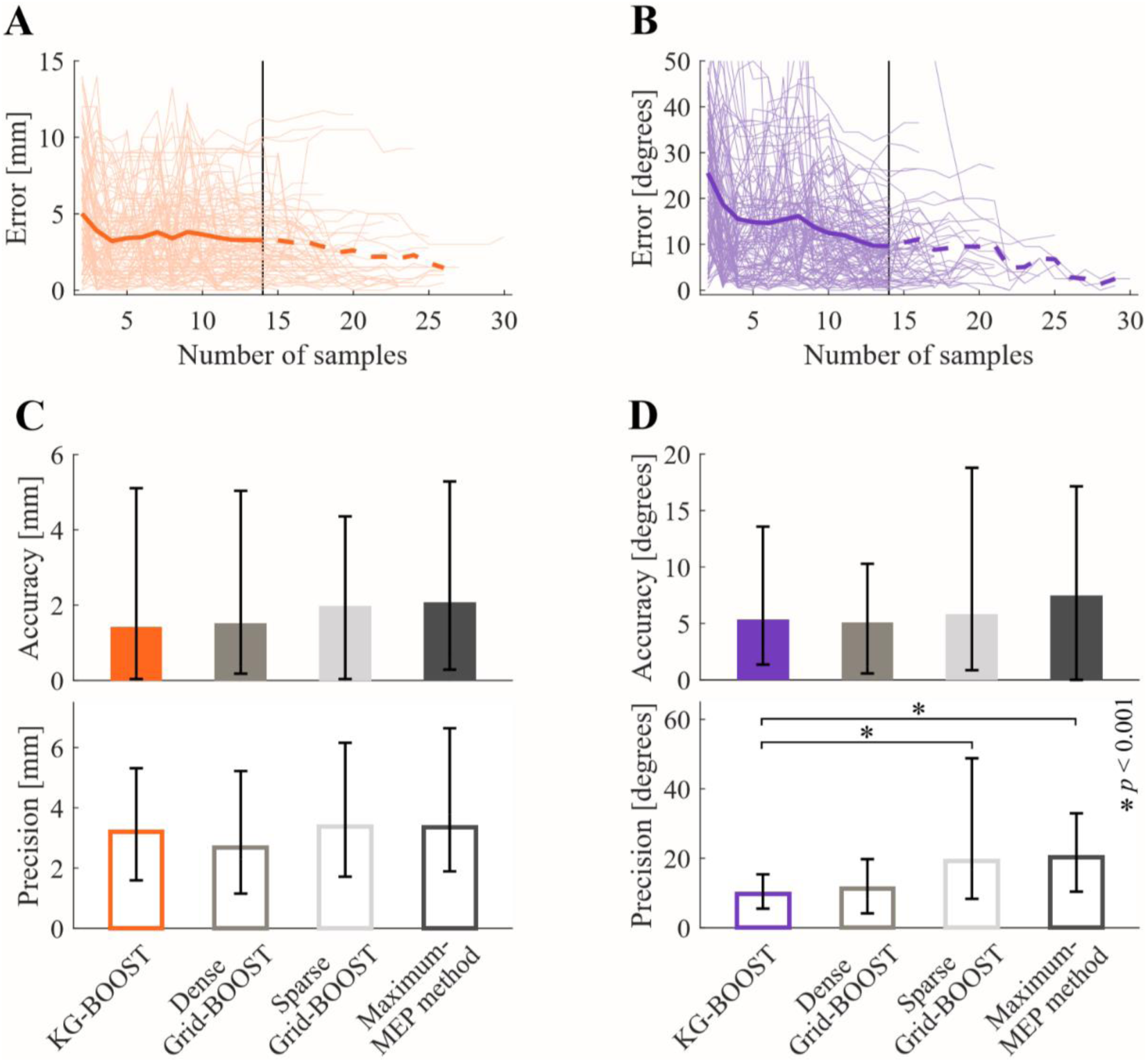
The convergence of KG-BOOST and the performance of the different search methods. A–B: The convergence of KG-BOOST in the search for the optimal stimulation location (A) and orientation (B). The light curves show how far from the ground truth the single runs are as a function of the number of samples. The solid dark curve depicts the average error from the ground truth until the minimum number of samples (14, black vertical line) has been reached. After 14 samples, as many of the runs have already finished, the average error curve (dashed line) includes only the remaining runs (until three or more left). C–D: Accuracy and precision of the different search methods in the search for the optimal stimulation location (C) and orientation (D). The bars depict the mean accuracy/precision of the search results with the black whiskers showing the minimum and maximum accuracy/precision over subjects and transducer placements. In D, the asterisks indicate the statistically significant differences in the precision of sparse Grid-BOOST and the maximum-MEP method compared to the precision of KG-BOOST.

The performance metrics of the orientation search can be found in Fig. 3D. The mean accuracy of KG-BOOST (mean: 5.4°; range 1.4–13.6°) was close to the ones of dense Grid-BOOST (mean: 5.1°; range: 0.6–10.3°; *p* = 0.82), sparse Grid-BOOST (mean: 5.8°; range: 0.9–18.8°; *p* = 0.78) and the maximum-MEP method (mean: 7.5°; range; 0–17.1°; *p* = 0.21). The mean precisions of KG-BOOST (mean: 9.7°; range 5.5–15.4°), dense Grid-BOOST (mean: 11.3°; range: 4.2–19.8°), sparse Grid-BOOST (mean: 19.1°; range: 8.3–48.8°) and the maximum-MEP method (mean: 20.2°; range: 10.4–32.9°) varied from each other. We found statistically significant differences in precision between KG-BOOST and both sparse Grid-BOOST (*p* = 0.000031) and the maximum-MEP method (*p* = 0.000001). The difference in precision between KG-BOOST and dense Grid-BOOST was not statistically significant (*p* = 0.25). The average number of samples acquired in the orientation search was 16 (range: 14–30; average time: 1.3 min) for KG-BOOST, sparse Grid-BOOST, and the maximum-MEP method. With dense Grid-BOOST, the number of samples was always 31 (average time: 2.6 min). In addition, the reference origin of the transducer placement O1, which was set to be perpendicular to the global orientation of the precentral gyrus, varied on average 6.7° (range: 1–13°) from the ground truth.

Our version of the AutoHS algorithm yielded an average error of 1.5 mm (range: 0.04–4.6 mm) and 19.7° (range: 2.1–37.9°) with respect to the ground truth for the location and orientation search, respectively. These values are not directly comparable with the accuracy values presented in Figs. 3C and 3D, since only one iterative search could be simulated for each subject and coil placement with the data available. Therefore, the results of AutoHS are excluded from Figs. 2 and 3 and the statistical analysis. With the AutoHS algorithm, the number of responses required for convergence was on average 53 (range: 20–110) and 62 (range: 20–163) for the location and orientation search, respectively, being about three to four times the number of stimuli used by KG-BOOST.

## 4 Discussion

We demonstrated that multi-locus TMS and Bayesian optimization can be successfully combined into an automated search of TMS targets. In this context, mTMS enables adjusting the stimulation location and orientation in a closed-loop setting without the need to move a coil, which significantly reduces the laboriousness of TMS. Bayesian optimization provides means to model and guide the stimulation in an effective and user-independent manner.

### 4.1 Performance of the automated target search

The automated online searches and further offline comparisons revealed that the mean accuracy in the location search was almost the same with all three versions of the BOOST algorithm (KG-BOOST guided with knowledge gradient and Grid-BOOST with dense and sparse sampling grids), when choosing the optimal target based on the maximal MEP response only, and with the AutoHS method (Harquel *et al.* 2017) (accuracies in the range of 1.4–2.1 mm). In the orientation search, the accuracy of the AutoHS method (19.7°) was worse than that of the other methods (5.1–7.5°). The small accuracy values indicate that the search results were centered almost symmetrically around the ground truth, which is expected behavior for any sensible search method.

The average precision of the location search was similar (2.7– 3.4 mm) among the four search strategies for which we were able to compute the precision (for AutoHS, we did not have enough samples for retrieving several independent search results). Thus, there were no statistically significant differences in precision in the location search between KG-BOOST and the other search methods. This can be explained by the fact that the motor maps extended over a large portion of the 30-mm-long search space and the median curves were relatively flat around their maximum (Fig. 2A). Instead, we found differences in the average precision of the optimal orientation search. KG-BOOST and dense Grid-BOOST had similar precisions (9.7° and 11.3°, respectively), while the precision of sparse Grid-BOOST and the maximum-MEP method were significantly worse (19.1° and 20.2°, respectively). These differences in the degree of scatter are not surprising, since the MEP-positive part of the median curve in the search space appeared narrower in the orientation search compared to the corresponding part in the location search. In the orientation search, KG-BOOST and dense Grid-BOOST got enough samples from the maximum area and the slopes of the mean MEP curve whereas sparse Grid-BOOST and the maximum-MEP method got fewer responses around the maximum, leading to larger deviation and, thus, worse precision in the search results.

Considering the efficiency of different search strategies, dense Grid-BOOST always used 31 samples, and AutoHS needed on average 53 and 62 samples, whereas the other methods used 18 and 16 samples on average in the location and orientation search, respectively. Sampling with KG-BOOST resulted in accuracy and precision similar to those of dense Grid-BOOST, but with approximately half of the number of samples. Also, the accuracies were similar (location search) or worse (orientation search) with AutoHS than with KG-BOOST, and AutoHS needed on average three to four times more samples for convergence. These findings indicate that KG-BOOST was more efficient than AutoHS and dense Grid-BOOST, and that sampling in a dense evenly spaced grid wastes samples especially in areas that produce no MEP responses. Results of the orientation search show that adaptive sampling with KG-BOOST led to better precision than placing the same number of samples evenly in the search space. Based on these results, we suggest using intelligent sampling, such as sampling with knowledge gradient (Frazier *et al.* 2009) that we used in KG-BOOST. This would allow one to avoid gathering too much data in areas with no MEPs while sampling adaptively around the maximum of the MEP curve to efficiently get enough information about the optimal target. We anticipate that even larger differences between the sampling methods are expected with larger or higher-dimensional search spaces (such as a two-dimensional location grid with additional variation in orientation), when non-guided grid sampling becomes very time-consuming.

Although the accuracy and precision values are generally good for almost all the search strategies, a single search outcome may still be several millimeters or degrees off from the ground truth regardless of the search method (see examples in Fig. 2). This is mainly due to the unavoidable high variability of the MEP responses that is present also in manual searches. Setting more strict stopping criteria would likely increase the accuracy and precision of KG-BOOST with the trade-off of increasing the number of samples needed and, thus, the measurement time. The stopping criteria can be chosen based on the needed accuracy and precision, which may differ between applications of the method.

### 4.2 Gaussian processes in target optimization

This study also demonstrated that Gaussian process regression is suitable for modeling the MEP response function (*i.e.*, motor map). Gaussian processes allow taking into account the uncertainties of the problem, the biggest source of uncertainty being the MEP variability. Another advantage of Gaussian process regression is its suitability for modeling a response function of any smooth shape as opposed to, *e.g.*, parametric Gaussian curve fitting, which assumes that the underlying function is a symmetric Gaussian distribution as in Harquel *et al.* (2017). We consider nonparametric fitting of MEP curves advantageous, since the motor maps can be asymmetric (see example maps in Weiss *et al*. 2013, Julkunen 2014, and van de Ruit *et al*. 2015). Sampling in a dense grid, as we did, is beneficial, since it assists in revealing the shape and, thus, the location of the peak of the response curve better than sampling in a coarse grid. Furthermore, as the Gaussian process regression model links the data of neighboring points, the expected convergence speed of a sampling in a dense grid is similar to that of a sampling in a sparse grid.

When modeling with Gaussian processes, one needs to ensure that the responses are handled on a suitable scale. We chose a logarithmic scale to satisfy better the assumption of location/orientation-independent MEP variance included in the model. The BOOST algorithm seems to tolerate well the variability in the MEP variance that occurs in practice. Note that on a logarithmic scale, the peaks of the MEP response curves (Fig. 2) appear broader than on a linear scale. For ensuring a sufficient number of samples and for placing them optimally, we suggest adaptively guiding the sampling with, *e.g.*, knowledge gradient (as in KG-BOOST) or entropy-based methods (as implemented by Harquel *et al*. (2017)). There are also other methods that could suit for efficient guiding of the sampling, such as the expected-improvement method (Mockus *et al*. 1978) or the use of confidence bounds as sampling criteria (Cox and John 1992), but these were not tested in this study.

Even though Gaussian process regression is a nonparametric method, it includes several model parameters that need to be tuned case-specifically. Our suggestion for determining the parameters in the case of TMS-target optimization are presented in Section 2.1.2, but there are also other ways to determine these parameters. To adapt the posterior adequately to the data while avoiding overfitting, it was crucial to correctly tune the smoothness parameter *a*_1,*d*_ (see Eq. (4)) to be of a reasonable magnitude. We chose to set this parameter with the simple zero-crossings formula and to keep it constant during the whole optimization procedure. Another option would be to tune *a*_1,*d*_ among the other model parameters adaptively and to teach the model with the acquired data. For this purpose, we also tested maximum-likelihood estimation and cross-validation (see Chapter 5 of Rasmussen and Williams 2005). However, in our experience, these approaches tended to overestimate *a*_1,*d*_, leading to overfitting the model to the data.

### 4.3 Future development and applications

Automated stimulation targeting could be further extended to adaptively adjust also the stimulation intensity (here, we used a predefined intensity, 110% rMT). In this case, the motor responses gathered during the target optimization might also be used in the rMT estimation, or the rMT could be determined separately after the target optimization with, *e.g.*, the adaptive algorithm presented by Awiszus (2003). In the future, the KG-BOOST algorithm can be applied for multi-dimensional problems, *e.g.*, to optimize simultaneously the location and orientation of the E-field, for example, with a 5-coil mTMS system similar to the one depicted by Koponen *et al*. (2018a). In principle, KG-BOOST can be implemented with any TMS system that allows automatic adjustment of the stimulation location and orientation, *e.g.*, with a robotically controlled stimulator. Furthermore, with a suitable software implementation, the algorithm could even guide the manual target search performed with a conventional navigated TMS system. With a multi-dimensional search space and a larger coverage of the cortex, one will be able to avoid the initial manual search, which we needed to place the transducer appropriately, and identify the optimal target even if it were situated abnormally.

The benefits of automated target optimization are applicable in several settings, from basic research to therapeutic uses of TMS. The automated target optimization could be used, *e.g.*, for studying the plasticity of motor cortex in a user-independent way. After modifying the function that guides the sampling, the BOOST algorithm may find applications in efficient and automated mapping of motor areas. If the actual cortical activation sites are of interest, one may combine the BOOST algorithm with individualized E-field modeling. In addition to finding the optimal stimulation parameters based on MEP amplitudes, the BOOST algorithm could be used to find an optimal stimulation target with respect to other available measures. For example, TMS targeting outside the motor cortex could be automatically optimized with respect to evoked cortical activity measured by electro-encephalography (Tremblay *et al*. 2019).

## 5 Conclusion

We demonstrated that electronically adjusted multi-locus TMS and Bayesian optimization provide a valid basis for automated search of TMS targets. The presented adaptively guided target search (KG-BOOST) gave results with good accuracy and precision, needing only a relatively small number of stimuli for convergence. We conclude that KG-BOOST enables fast, easy and user-independent target optimization, and that its benefits are applicable from basic research to therapeutic uses of TMS.

## Acknowledgements

We thank Emma Skarstein for contributing to the literature search and investigating different aspects of modeling with Gaussian process regression. We thank Selja Vaalto and Ulf Ziemann for useful comments on the manuscript. This project has received funding from the Academy of Finland (Decisions No. 294625, 306845, and 327326), the Finnish Cultural Foundation, and the European Research Council (ERC) under the European Union’s Horizon 2020 research and innovation programme (grant agreement No 810377).

## Appendix

Here, we present further details for computing the knowledge gradient. Assume that we have *N* noisy samples **y** = [*y*_1_, …, *y*_*N*_]^T^ that correspond to the sampling parameters **X** = [**x**_1_, …, **x**_*N*_]. Knowing them, we choose the next sampling parameters **x**_*N*+1_ with a knowledge-gradient sampling policy (Frazier *et al*. 2009, Frazier and Wang 2015). For getting the maximum of KG(**x**) (approximately), we choose an appropriately dense subset 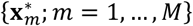 in the search space. For a fixed 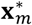, we form sequences *A*(*r*) and *B*(*r*), *r* = 1, …, *M*, by setting for each 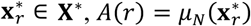 as in Eq. (6) and

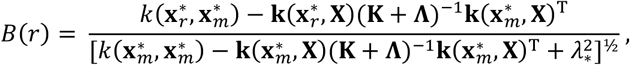

where 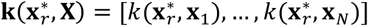 and 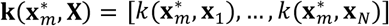 contain the covariances between 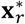 or 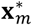 and **x**_1_, …, **x** _*N*_ (formula for *k* in Eq. (4)), **Λ** is a diagonal matrix with 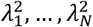 on its diagonal, and 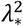 is the noise variance at location 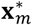. As in Frazier *et al*. (2009), we choose subsequences *α*(*s*) and *β*(*s*) of *A*(*r*) and *B*(*r*) and an additional sequence *γ*(*s*), *s* = 1, …, *S*, by Matlab codes *AffinebreakpointsPrep.m* and *Affinebreakpoints.m* given in Frazier (2010). With these sequences, we get the knowledge-gradient function as

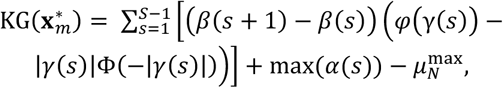

where 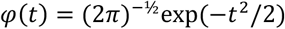 and 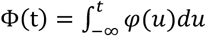 are the probability density function and the cumulative density function of a normalized Gaussian random variable, respectively. Finally, we get the next sampling point as

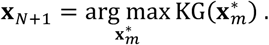

